# Insights into the molecular mechanism and management of resistant *Echinochloa crus-galli* biotypes from the Philippines to acetolactate synthase inhibitor bispyribac-sodium

**DOI:** 10.1101/2025.10.03.680269

**Authors:** Niña Gracel Dimaano, John Kenneth Polo, Euzane Joy Macarilay, Christianne Lei Golena, Micah Rinoa Testor, Jessa Jean Buen, Clare Hazel Tabernilla, Analiza Henedina Ramirez, Mary Joy Abit, Satoshi Iwakami

## Abstract

Rice production in the Philippines is heavily reliant on herbicide application for weed control. Acetolactate synthase (ALS) inhibitors are the mostly used herbicides for combating major weeds associated to rice, such as *Echinochloa crus-galli*. In separate locations from the Philippines, rice farmers expressed concerns that the application of ALS inhibitor bispyribac-sodium has been showing poor control of *E. crus-galli*. Here, we confirmed and quantified the level of bispyribac-sodium resistance in the putative resistant biotypes. Resistance was confirmed in R4 and R12 biotypes collected from the northern and central parts of the Philippines, respectively. The estimated GR_50_ (dose resulting in 50% growth reduction) revealed a 36.7-and 60.3-fold resistance in R4 and R12 biotypes (106.8 g a.i. ha^-1^ and 175.4 g a.i. ha^-1^) compared to the S biotype (2.91 g a.i. ha^-1^). The R4 biotype carries a single nucleotide mutation in the *ALS2* gene translating to Trp-574-Leu amino acid substitution, while there was no mutation detected in any of the *ALS* conserved regions of R12. Pre-treatment of malathion increased the sensitivity of R12 to bispyribac-sodium, suggesting the involvement of cytochrome P450s. Sensitivity assays to alternative post-emergence herbicides showed that both R4 and R12 can be effectively controlled by metamifop [acetyl-CoA carboxylase (ACCase) inhibitor]. However, R12 survived the application of fenoxaprop-*P*-ethyl + ethoxysulfuron (ACCase inhibitor + ALS inhibitor). This study unravels the molecular mechanism of the first resistance case to ALS inhibitor in the Philippines and provide practical implications for efficient herbicide resistance management in the affected rice areas.

## 1. Introduction

Herbicides have revolutionized agriculture by rendering a reliable, efficient, and economical approach to control weeds. The effective weed control provided by herbicides has greatly contributed to a significant increase in crop yields (Heap, 2014). However, the dramatic increase in usage and reliance on herbicides has created strong selection pressure in weed populations that has allowed rare resistance genes to be selected, enriched, and result in the rapid evolution of herbicide resistance in weed populations (Powles and Yu, 2014). The rapid increase in the cases of herbicide resistance in weeds combined with the lack of herbicides with novel modes of action (MOA) threatens crop production worldwide (Heap, 2014).

The mechanism of herbicide resistance in weeds can be explained by either (1) target-site resistance (TSR) endowed by mutations or amplification of herbicide target protein, or (2) non-target-site resistance (NTSR) conferred by the activity of herbicide-metabolizing enzymes, reduced herbicide uptake or translocation, or other mechanisms not involving the herbicide targets (Yu and Powles, 2014). Elucidation of the mechanism(s) conferring herbicide resistance is necessary for a better understanding of the rapid evolution of resistance in weed species, identifying effective herbicide options, and designing better herbicide management plans to successfully control and mitigate the spread of the herbicide resistant populations.

Acetolactate synthase (ALS) inhibitor herbicides are among the frequently used herbicides in agricultural fields since its introduction to the market in the early 1980s (Tranel and Wright, 2002; Zhou et al., 2007). ALS inhibitors inhibit the catalytic activity of ALS, the first enzyme in the biosynthesis of branched-chain amino acids, leading to the disruption of the synthesis of valine, leucine, and isoleucine (Lonhienne et al., 2022). Due to the long history of the availability of ALS inhibitors, many weed species have evolved resistance to this herbicide group (Tranel and Wright, 2002). Currently, there are a total of 176 reported resistance cases (108 in dicots and 68 in monocots) for ALS inhibitors (Heap, 2025).

*Echinochloa crus-galli* (L.) Beauv. (barnyardgrass) is considered a serious weed in rice production systems in many countries worldwide (Damalas, 2008). It is an allohexaploid species (2n = 6× = 54) that is assumed to have developed from the hybridization between the tetraploid species *Echinochloa oryzicola* (2n = 4× = 36) and an unknown diploid species (2n = 2× = 18) (Ye et al., 2020). *Echinochloa crus-galli* shows morphological and physiological adaptations that allow it to be highly competitive with rice (Chauhan and Johnson, 2010). Season-long *E. crus-galli* interference has been reported to reduce grain yields of rice cultivars by 28 to 65% (Smith, 1988). *Echinochloa crus-galli* is among the weed species with several confirmed resistance cases to ALS inhibitors. To date, 15 countries have reported cases of resistance to ALS inhibitors in *E. crus-galli* populations (Heap, 2025).

In the recent years, farmers in the Philippines have expressed poor control of *E. crus-galli* in their rice fields despite the application of the recommended rates of ALS inhibitors.

The existence of resistant *E. crus-galli* biotypes to ALS inhibitors is of particular concern since the rice production systems in the country are heavily reliant on the limited herbicide options that can be selectively applied to rice. Here, we have confirmed the resistance to ALS inhibitor bispyribac-sodium in the putative resistant *E. crus-galli* biotypes and provided substantial evidence on the molecular mechanism endowing the resistance. The elucidation of the resistance mechanisms will contribute to modeling effective management strategies and mitigation measures to prevent the further spread of resistant biotypes.

## 2. Materials and Methods

### 2.1 Origin and Preparation of Plant Materials

The seeds of putative resistant (R) biotypes of *E. crus-galli* that survived the recommended rate of ALS inhibitor bispyribac-sodium were collected in bulk (20 plants per approximately 2500 m^2^ field) from five sites in the Philippines (Table 1). The sites with suspected R biotypes were selected based on persistent complaints of rice farmers regarding uncontrolled *E. crus-galli* in their fields and herbicide application history (continuous use of bispyribac-sodium for at least five years). The susceptible (S) *E. crus-galli* biotype was collected within the vicinity of the University of the Philippines Los Baños (UPLB), Laguna, Philippines where no history of bispyribac-sodium was applied.

**Table 1.** Sensitive and putative resistant biotypes of *Echinochloa crus-galli* used in the initial herbicide sensitivity screening to acetolactate synthase (ALS) inhibitor, bispyribac-sodium.

| Biotype ID | Place of collection |  | Sensitivity to bispyribac-sodium |
| --- | --- | --- | --- |
|  | Province | Municipality |  |
| 4 | Nueva Vizcaya | Bayombong | Survived |
| 5 | Pangasinan | Urdaneta | Complete control |
| 10 | Bulacan | San Rafael | Complete control |
| 12 | Bulacan | Baliuag | Survived |
| 15 | Pampanga | San Luis | Complete control |
| S | Laguna | Los Baños | Complete control |

Note: S biotype, previously confirmed to be sensitive to bispyribac-sodium. The recommended rate of bispyribac-sodium was applied to the *E. crus-galli* biotypes at 10 days after sowing. The biotypes that survived at 14 days after treatment were transferred to bigger pots for self-propagation and seed purification prior to dose-response assays.

### 2.2 Sensitivity Assays to Bispyribac-sodium

Preliminary sensitivity assays were conducted on the S and putative R biotypes of *E. crus-galli.* Seeds were sterilized according to the procedure of Dimaano et al. (2020) and pre-germinated in Petri plates lined with moistened filter paper at 30°C with a 12-hour photoperiod. Twelve pre-germinated seeds per each biotype were then transferred into five sets of pots (20 cm x 25 cm) filled with sterilized soil then added with complete NPK fertilizer (14-14-14). Plants were maintained in a shallow flooded condition before herbicide application. At 10 days after sowing, the seedlings were sprayed with the recommended rate of bispyribac-sodium (Nominee SC 100, 32 g a.i. ha^-1^, Bayer Crop Science, Inc.) using a knapsack sprayer calibrated to deliver 160 L ha^-1^. A non-ionic surfactant (polyethylene sorbitan acid and polyoxyethylene dodecyl ether; Agrisol 150K, 45 g a.i. ha^-1^, Kao Corporation, Tokyo, Japan) was added in accordance with the label recommendations. The sensitivity of the putative R biotypes was compared to that of the S biotype. The seedlings that survived after bispyribac-sodium application were individually transferred into bigger pots and grown for self-propagation and seed purification prior to dose-response assays.

To confirm and quantify the level of resistance to bispyribac-sodium, dose-response assays on the S and putative R biotypes were performed in a screenhouse at UPLB from July to September of 2022. Each R biotype used was from the next-generation purified seeds of a single plant that survived the preliminary sensitivity assay. Seed and soil preparations were the same as described above. Five to eight seedlings of each biotype were transplanted into 20 cm x 25 cm pots and thinned out to three plants per pot at one week after transplanting. Plants at three-to four-leaf collar stages were treated with 0, 8, 16, 24, 32, 64, 96, 128, 192, 256, and 288 g a.i. ha^-1^ of the commercial formulation of bispyribac-sodium (Nominee SC 100, Bayer Crop Science, Inc.) added with a non-ionic surfactant (same as previous section). Control plants were treated with distilled water added with surfactant. Herbicide treatments were applied using a CO_2_-pressurized backpack sprayer equipped with 8004 flat fan nozzle and calibrated to deliver 160 L ha^-1^. Each experiment was conducted at least twice and replicated three times. At 21 DAT, the responses of S and putative R biotypes were visually assessed, and the above ground biomass was harvested and oven-dried at 60°C for three days. The dry weight data were measured and expressed as percentage of the mean of untreated plants. The 50% inhibitory dose (GR_50_) for each biotype was estimated by log-logistic regression with the ‘drc’ package (Ritz et al., 2015) in R version 3.4.4 (R Core Team, 2018). Resistance index (RI) was computed as GR_50_ of the R biotype over GR_50_ of the S biotype.

### 2.3. Sensitivity to Alternative Post-emergent Herbicides

To determine the bioefficacy of alternative post-emergent herbicides for the effective management of R biotypes, the recommended rates of the commercial formulations of ACCase inhibitor metamifop (Pyzero 10 EC, 75 g a.i. ha^-1^, FMC Corporation) and ACCase inhibitor + ALS inhibitor fenoxaprop-*P*-ethyl + ethoxysulfuron (Ricestar Xtra 89 OD, 34.5 g a.i. ha^-1^ + 10.5 g a.i. ha^-1^, Bayer Crop Science, Inc.) were applied to the S and R biotypes at four-leaf collar stage. The surfactant added in the herbicide treatments as well as the sprayer used in the assays were the same as described above. The sensitivity of each R biotype to the herbicides was compared to that of the S biotype. Each experiment was conducted at least twice and replicated three times.

### 2.4. Sensitivity Assay to Bispyribac-sodium after Malathion Treatment

To determine the probable involvement of cytochrome P450s in the resistance mechanism, a known cytochrome P450 inhibitor, malathion (Malathion 57 EC, 1000 g a.i. ha^-1^, Vast Agro Solutions Inc.), was sprayed one hour before the application of the varying rates of bispyribac-sodium (0, 32, 64, 96, and 128 g a.i. ha^-1^). Each experiment was conducted at least twice and replicated three times. At 21 DAT, the above ground biomass was harvested and oven-dried at 60°C for three days for collection of dry weight data. The GR_50_ for each biotype was estimated by non-linear four-parameter log-logistic model described by Seefeldt et al. (1995) using the Sigma Plot 14.5 (SPSS IBM® Corporation, Route 100, Somers, NY).

### 2.5 DNA Extraction and Gene Sequence Analysis

DNA was extracted from the young leaves of each biotype of *E. crus-galli* seedlings using the GF-1 Plant DNA Extraction kit (Vivantis, Malaysia) following its protocol. The seedlings were grown from self-pollinated single plants of the R biotypes that survived the preliminary herbicide sensitivity assays. The three copies of *ALS* genes were amplified using the referenced primer sets and PCR protocols from the study of Iwakami et al. (2015). The PCR products were treated with ExoSAP-IT (Thermo Fisher Scientific, USA) and directly sequenced. The resulting sequence fragments were analyzed using Bioedit Sequence Alignment Editor 7.2 (Hall, 2011).

## 3. Results

### 3.1 Sensitivity of putative resistant *E. crus-galli* biotypes to bispyribac-sodium

Based on preliminary bispyribac-sodium sensitivity assays, only the putative R biotypes from Bayombong, Nueva Vizcaya (biotype 4, R4) and Baliuag, Bulacan (biotype 12, R12) survived the application of the recommended rate of bispyribac-sodium. The biotypes S, 5, 10, and 15 had suppressed growth and chlorosis at as early as 7 days after treatment (DAT). All tested biotypes except R4 and R12 turned brown and eventually died after two weeks of herbicide application (Table 1). No segregation was observed in the herbicide sensitivity of all the seedlings of each biotype. Subsequently, R4 and R12 biotypes that survived were transferred into bigger pots for self-propagation and seed purification. Then, the purified seeds were subjected to dose-response assays to confirm and quantify the resistance level and to elucidate the molecular mechanism of the first resistance case to ALS inhibitor in the Philippines.

Dose response assays showed that the recommended rate of bispyribac-sodium (32 g a.i. ha^-1^) caused visible injury on the S biotype. Leaf chlorosis on the S biotype was first observed at 5 DAT. By 7 DAT, complete yellowing of the leaves was evident which progressed to plant necrosis, while at 14 to 21 DAT, complete control of the S biotype plants was observed. On the contrary, the putative R biotypes showed little injury even after the application of higher doses of bispyribac-sodium (Figure 1). Resistance to bispyribac-sodium was confirmed by the GR_50_ parameters computed for biotypes R4 and R12. The estimated GR_50_ revealed a 36.7-and 60.3-fold bispyribac-sodium resistance in R4 and R12 biotypes (106.8 g a.i. ha^-1^ and 175.4 g a.i. ha^-1^) compared to the S biotype (2.91 g a.i. ha^-1^).

**Figure 1.**
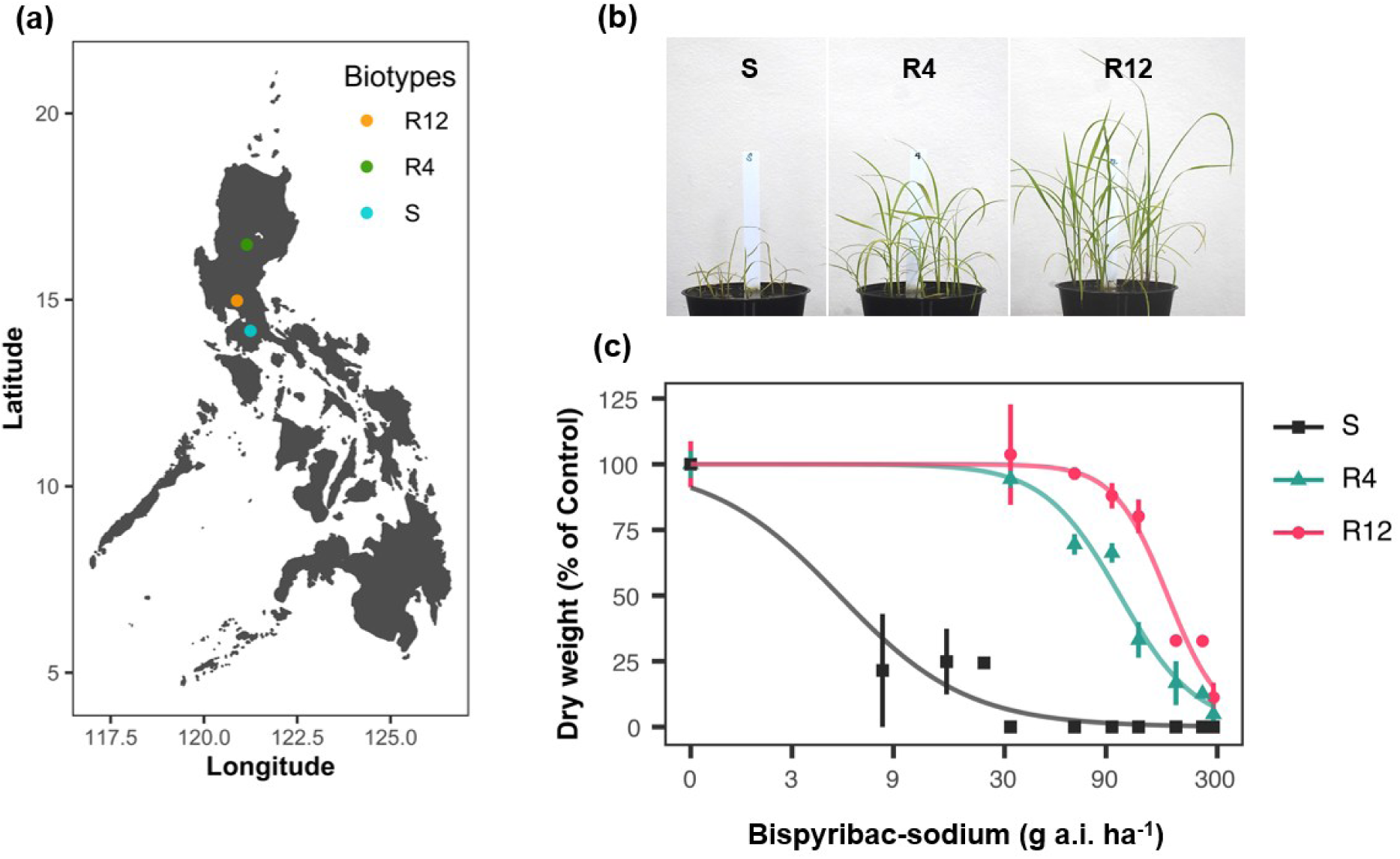
Herbicide sensitivity of susceptible (S) and resistant (R) biotypes of *Echinochloa crus-galli* to bispyribac-sodium. (a) Collection sites of the S, R4, and R12 biotypes corresponding to Laguna, Nueva Vizcaya, and Bulacan, Philippines, respectively. (b) Sensitivity of the S and R biotypes to the recommended field rate of bispyribac-sodium at as early as 7 days after treatment. (c) Dose response curve of S and R biotypes to the varying concentrations of bispyribac-sodium.

### 3.2 Sensitivity of bispyribac-sodium resistant biotypes to alternative herbicides

Alternative post-emergence herbicides were evaluated for probable effective control of the R biotypes. Results showed that both R4 and R12 biotypes were effectively controlled by metamifop, an ACCase inhibitor, when applied at the recommended dose (75 g a.i. ha^-1^). Similar phytotoxicity symptoms were observed in both S and R biotypes at 21 DAT (Figure 2). Another post-emergence herbicide, fenoxaprop-*P*-ethyl + ethoxysulfuron (ACCase inhibitor + ALS inhibitor) (34.5 g a.i. ha^-1^ + 10.5 g a.i. ha^-1^) showed phytotoxicity against the R4 biotype; however, the R12 biotype exhibited resistance to the herbicide. The herbicide phytotoxicity symptoms observed on the R4 biotype were similar to that on the S biotype, but the R12 biotype survived and produced green healthy leaves and productive tillers despite the application of the herbicide recommended rate (Figure 2).

**Figure 2.**
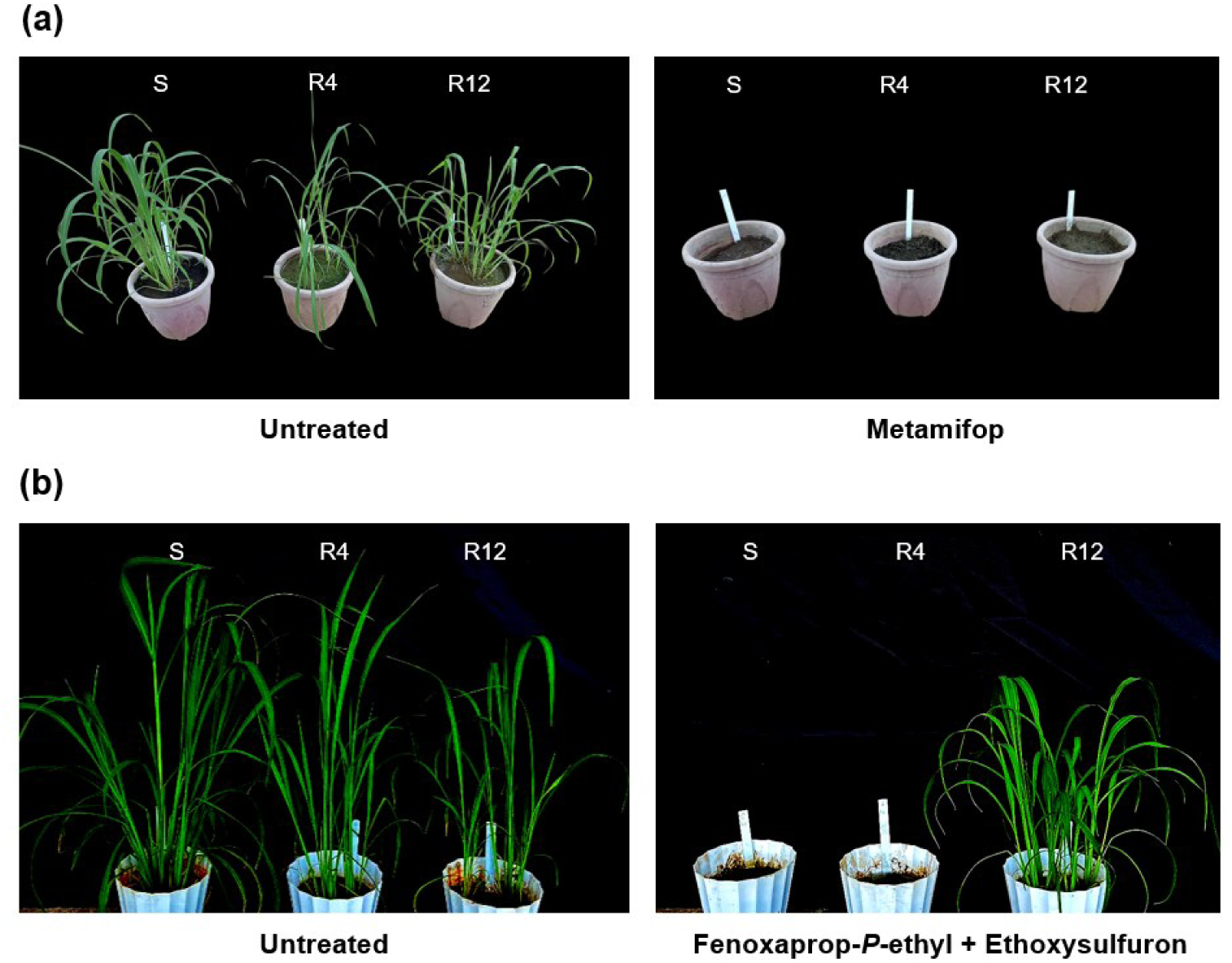
Herbicide sensitivity of susceptible (S) and resistant (R) biotypes of *Echinochloa crus-galli* to the recommended rates of metamifop (75 g a.i. ha^-1^) and fenoxaprop-*P*-ethyl + ethoxysulfuron (34.5 g a.i. ha^-1^ + 10.5 g a.i. ha^-1^).

### 3.3 Sequencing of herbicide target-site genes

The three copies of *ALS* genes (Iwakami et al., 2015) of the two resistant *E. crus-galli* biotypes, R4 and R12, were analyzed to confirm if the underlying herbicide resistance mechanism is TSR-based. A homozygous single-nucleotide substitution of G to T, translating to Trp-574-Leu change in amino acid sequence, was detected in one of the *ALS* genes of the R4 biotype (Figure 3), a gene located in subgenome A in *E. crus-galli* genome (Wu et al., 2022), which was previously named as *ALS2* (Iwakami et al., 2015). No known single nucleotide polymorphism (SNP) or mutation endowing resistance was detected in the other two *ALS* gene sequences of R4. Similarly, no SNPs or mutations were detected in all the *ALS* gene sequences of the R12 biotype.

**Figure 3.**
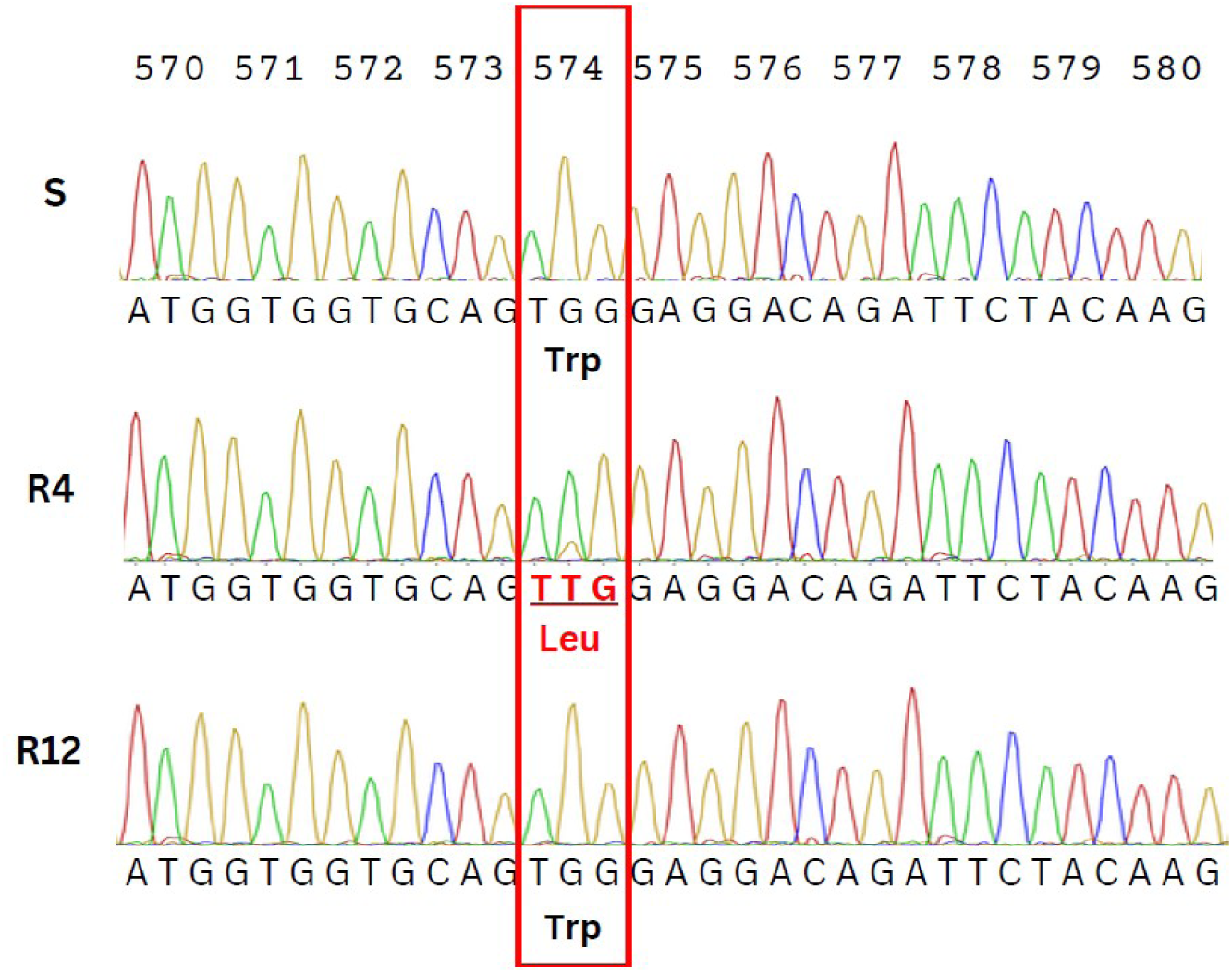
Alignment of partial *ALS2* gene sequences of susceptible (S) and resistant biotypes (R4 and R12) showing the Trp-574-Leu mutation that confers resistance to acetolactate synthase inhibitor, bispyribac-sodium.

The sensitivity testing of R12 to ACCase inhibitor + ALS inhibitor (fenoxaprop-*P*-ethyl + ethoxysulfuron) resulted in the survival of the biotype after herbicide application; thus, the *ACCase* gene sequences of R12 were analyzed to check whether the mechanism of resistance is also TSR-based. Based on the gene sequence analysis, no mutation endowing resistance to ACCase inhibitors was found in all the six *ACCase* genes (Iwakami et al., 2024) of the R12 biotype.

### 3.4 Investigation of the involvement of P450-mediated resistance mechanism

Since no mutation was found in the *ALS* genes of the R12 biotype, it was suspected that the mechanism of resistance is NTSR-based, particularly due to the metabolizing activity of herbicide detoxifying enzymes such as cytochrome P450s (P450s). Here, the effect of malathion, a known P450 inhibitor, was evaluated to provide evidence that the mechanism of resistance in the R12 biotype is possibly conferred by the metabolizing activity of P450s. Results showed that the application of malathion caused an increased sensitivity of R12 biotype to bispyribac-sodium. The pre-treatment of malathion decreased the GR_50_ (rate to cause 50% growth reduction) in the R12 biotype from 193.53 g a.i. ha^-1^ to 51.98 g a.i. ha^-1^ (Table 2).

**Table 2.**
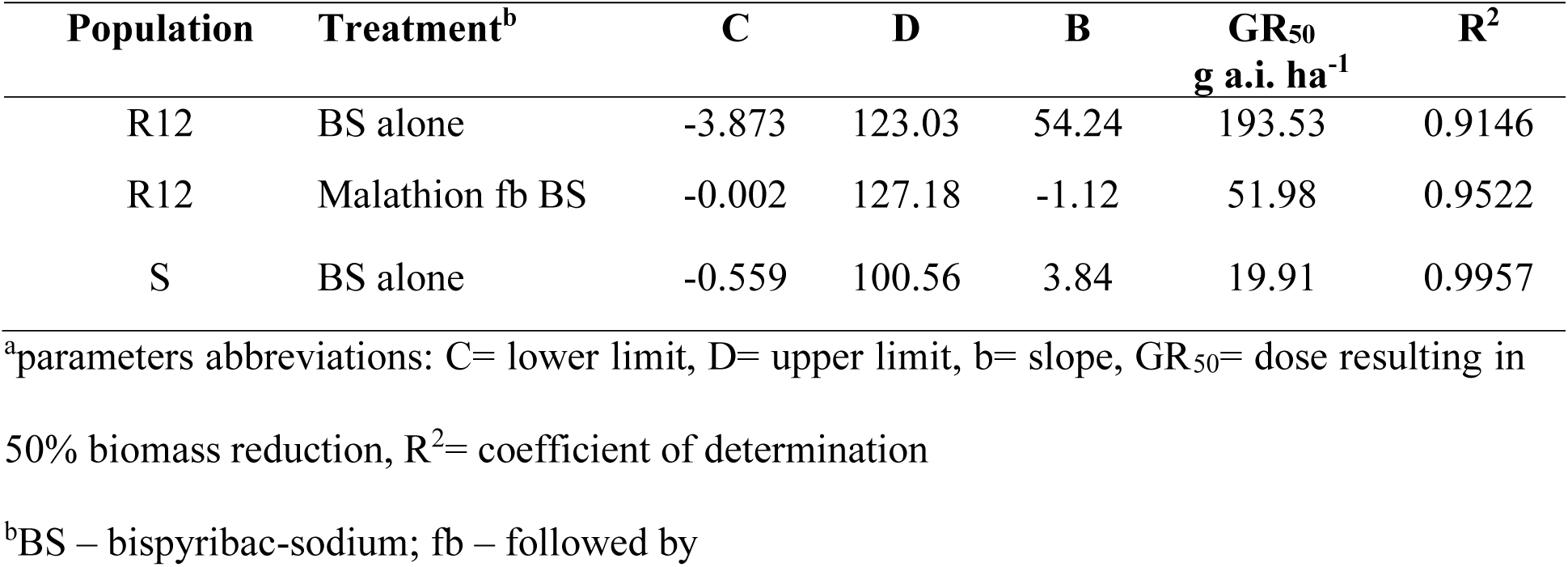
Log-logistic parameters^a^ of *Echinochloa crus-galli* biomass reduction and 50% inhibitory dose (GR_50_) at 21 days after treatment (DAT).

| Population | Treatment<sup>b</sup> | C | D | B | GR<sub>50</sub><br>g a.i. ha<sup>-1</sup> | R<sup>2</sup> |
| --- | --- | --- | --- | --- | --- | --- |
| R12 | BS alone | -3.873 | 123.03 | 54.24 | 193.53 | 0.9146 |
| R12 | Malathion fb BS | -0.002 | 127.18 | -1.12 | 51.98 | 0.9522 |
| S | BS alone | -0.559 | 100.56 | 3.84 | 19.91 | 0.9957 |

<sup>a</sup>parameters abbreviations: C= lower limit, D= upper limit, b= slope, GR<sub>50</sub>= dose resulting in 50% biomass reduction, R<sup>2</sup>= coefficient of determination
<sup>b</sup>BS – bispyribac-sodium; fb – followed by

## 4. Discussion

### 4.1 Confirmed resistance in *E. crus-galli* biotypes

The pressing issue of poor herbicide control of *E. crus-galli* in selected areas in the Philippines was confirmed to be due to herbicide resistance in two distinct biotypes, R4 and R12. Molecular analysis revealed that the two biotypes exhibit distinct resistance mechanisms. The R4 biotype showed target-site based resistance solely to the ALS inhibitor bispyribac-sodium, while the R12 biotype does not carry any mutation in the ALS target site gene. This suggests that the mechanism of resistance in the R12 biotype is most likely NTSR-based. Furthermore, the R12 biotype exhibited a concerning multiple-herbicide resistance to ALS and ACCase inhibitors, as the biotype showed no sensitivity to the application of fenoxaprop-*P*-ethyl + ethoxysulfuron. This finding underscores the urgent need for further elucidation of the resistance mechanism in the R12 biotype, and formulation of suitable strategies to manage the resistance.

An intriguing aspect of resistance evolution in *E. crus-galli* is its hexaploid nature, resulting in a large number of herbicide target-site gene copies. The presence of multiple copies in herbicide target-site genes sometimes leads to distinctive patterns of resistance evolution such as biased dependency on specific copies for resistance evolution. For example, *Monochoria vaginalis*, a noxious paddy weed, employs only two copies of ALS genes, not the other three copies, in resistance evolution (Tanigaki et al., 2021). Similarly, biased preference to one ALS copy was also reported in the study of global *E. crus-galli* populations, where TSR mutation was preferably found in an *ALS* gene in subgenome A (Wu et al., 2022). The Trp-574-Leu mutation in the R4 biotype was also identified in the same *ALS* copy. It should be interesting to investigate the mechanism of biased representation of this *ALS* copy in the resistance evolution of *E. crus-galli*.

Trp574Leu mutation confers resistance to ALS inhibitors of all chemical classes (Yu and Powles, 2014). The cross-resistance of this particular mutation is due to the importance of Trp-574 binding site in defining the shape of ALS active-site channel as well as for anchoring ALS herbicides (McCourt et al., 2006; Powles and Yu, 2010). Although we have not tested it yet, the R4 biotype would most likely show resistance to other ALS herbicides from different chemical families.

The R12 biotype exhibited resistance to ALS herbicide bispyribac-sodium, although no TSR mutation was observed at the target site. Treatment with the P450 inhibitor malathion significantly increased sensitivity to bispyribac-sodium, suggesting P450 involvement in bispyribac-sodium resistance. Nevertheless, since the influence of malathion to bispyribac-sodium response was only analyzed in the R population, it is crucial to note that conclusive evidence regarding whether P450 inhibition by malathion explains the difference between R and S populations is lacking. Though we have not investigated it in the current study, the R4 biotype may also have underlying NTSR mechanism. A more in-dept investigation is necessary to unravel the probable NSTR mechanisms in both R4 and R12 biotypes.

P450s are versatile monooxygenases that catalyze a wide range of reactions and are encoded by hundreds of genes in plant genomes. Certain P450 enzymes can detoxify herbicides, playing a crucial role in the evolution of herbicide resistance in weeds (Dimaano and Iwakami, 2021). In *Echinochloa* spp., several P450 enzymes that detoxify ALS inhibitors have been identified from the CYP81A subfamily (Dimaano et al., 2020; Pan et al., 2022). However, no clear activity against bispyribac-sodium has been detected so far, suggesting the involvement of other P450 subfamily in the bispyribac-sodium resistance of R12. In rice, a gene from the CYP72A subfamily plays a major role in bispyribac-sodium metabolism (Saika et al., 2014). Similar genes in *E. crus-galli* may be involved in bispyribac-sodium resistance in the R12 biotype.

Notably, the R12 biotype survived the recommended rate of ACCase inhibitor + ALS inhibitor (fenoxaprop-*P*-ethyl + ethoxysulfuron). Since the application history of fenoxaprop-*P*-ethyl in the field where R12 was collected is unclear, it is unknown whether the evolution of resistance to bispyribac-sodium and fenoxaprop-*P*-ethyl is independent. On the other hand, research cases so far suggest that the resistance to the two herbicides is caused by metabolism-based cross-resistance: cross-resistance to ALS and ACCase inhibitors is often observed in metabolic resistant biotypes in *Echinochloa* spp. (Im et al., 2009; Iwakami et al., 2014, 2019, 2024; Pan et al., 2022). Although the detailed mechanism of the cross-resistance remains elusive, co-regulation of multiple enzymes with different activities towards herbicides is proposed (Suda et al., 2023). In the Poaceae family including *Echinochloa*, fenoxaprop-*P*-ethyl is detoxified via glutathione conjugation (Tal et al., 1993; Bakkali et al., 2007). If P450 is involved in bispyribac-sodium resistance in R12 as suggested from the malathion experiment, a mechanism that co-regulate P450 and glutathione S-transferase might underlie the multiple-herbicide resistance.

### 4.2 Insights into the management of the resistant biotypes in the Philippines

Weed control in Philippine rice fields is highly dependent on herbicides combined with handweeding operations (Gianessi and Williams, 2012). Selective chemical weed management in the country began in 1948 with the introduction of 2,4-D for broadleaved weed control. In the 1960s and 1970s, herbicides such as butachlor, propanil, and thiobencarb were quickly adopted by rice farmers for their effectiveness in grass control, followed by the introduction of oxadiazon, pretilachlor, bentazon, and pendimethalin in the 1970s and 1980s. Later, in the 1980s and 1990s, fenoxaprop-*P*-ethyl and cyhalofop-butyl, as well as sulfonylureas were introduced for post-emergence weed control, and by the late 1990s and early 2000s, penoxsulam and bispyribac-sodium became part of rice weed management practices (Casimero et al., 2023). Currently, various herbicides are available to control annual grasses, such as *Echinochloa* spp. Post-emergence herbicide options include ALS inhibitors (e.g., pyribenzoxim), ACCase inhibitors (e.g., metamifop), and photosystem II inhibitors (e.g., propanil). Pre-emergence herbicide options include very long chain fatty acid inhibitors (e.g., butachlor), PPO inhibitors (e.g., oxadiazon); and microtubule assembly inhibitors (e.g., pendimethalin). Additionally, deoxy-D-xylulose phosphate synthase inhibitor clomazone is effective as both pre-and early post-emergence herbicide (Fertilizer and Pesticide Authority, 2025).

Despite the range of pre-and post-emergence herbicide options, Filipino farmers hesitate to use pre-emergence herbicides due to their perception that these herbicides might cause phytotoxicity when applied on direct-seeded rice; preference to apply pre-emergence herbicide when weeds are visible (2 leaf-stage at the minimum); and option not to disturb the soil layer as they perceive that weeds will emerge when the soil is disturbed. Hence, farmers often rely on post-emergence herbicides, especially ALS inhibitors that are effective against *E. crus-galli* that have already emerged in their fields. Moreover, the rice areas where the resistant *E. crus-galli* biotypes were collected are with constrained irrigation sources, thus flooding the field as an alternative weed management tactic is not a viable option. Due to these farming practices and environmental constraints, farmers have become reliant on few ALS inhibitors for managing *E. crus-galli*.

The findings of this study emphasize the threat of TSR-based resistance (e.g., Trp574Leu mutation) to the efficacy of available ALS inhibitors in the country, as resistance can be extended to other chemical groups (e.g., imidazolinones, pyrimidinyl benzoates, triazolopyrimidines, and sulfonylureas). The other alternative options of farmers in controlling *E. crus-galli* include ALS inhibitors such as flucetosulfuron, halosulfuron-methyl, imazosulfuron, propyrisulfuron, penoxsulam, and pyribenzoxim (FPA, 2025). Even more threatening is that resistance can be further endowed to new ALS inhibitor herbicides that are yet to be introduced in the market.

Effective information dissemination is crucial to educate farmers about herbicide resistance mechanisms and development. Currently, many Filipino farmers have minimal to no knowledge about herbicide resistance. Providing comprehensive education on herbicide resistance can empower farmers to develop strategies to combat the resistance cases in their fields. As mentioned above, an effective approach to managing the resistant biotypes is to diversify the herbicide MOA, moving beyond the commonly available ALS inhibitors. The efficacy of pre-emergence herbicides, which farmers are hesitant to use, should be tested.

However, caution is necessary as resistance to pre-emergence herbicide (e.g., butachlor) has been previously reported in the country (Juliano et al., 2010). In addition to chemical weed control, implementing cultural weed control is also needed. Although flooding is not viable due to the constrained irrigation sources, other techniques such as the stale seedbed method, where weed seeds are eradicated before the next planting season, can be an effective option. To control herbicide-resistant weeds and minimize the evolution of future resistance, it is necessary to employ a diverse range of strategies for weed management.

## 5. Conclusion

In conclusion, this study confirmed and elucidated the mechanisms of the first case of *E. crus-galli* resistance to ALS inhibitor bispyribac-sodium in the Philippines. Sequence analysis of *ALS* genes confirmed the TSR mechanism in the R4 biotype conferred by Trp-574-Leu mutation, while the resistance in R12 could be metabolism-based. Further investigation of the suspected NTSR mechanism in R12 as well as the possible coexistence of both TSR and NTSR in R4 is crucial for developing effective management and preventive strategies for these resistance cases. Testing other herbicides with different MOA is necessary to determine the extent of cross-and multiple-resistance in the resistant biotypes. Furthermore, farmers’ surveys should be conducted to determine the magnitude of spread of resistance in their fields and their perception on the management of herbicide resistance.

## Acknowledgments

This research was funded by the UPLB Basic Research Program, Office of the Vice Chancellor for Research and Extension awarded to Niña Gracel Dimaano and by the JSPS KAKENHI grant JP22H02347 awarded to Satoshi Iwakami.

## Author Contributions

NG Dimaano, MJ Abit, and S Iwakami contributed to the research conception and design. NG Dimaano and CH Tabernilla collected and prepared the materials. All authors contributed to data collection, analysis, interpretation, and discussion. All authors contributed to the manuscript preparations. NG Dimaano, MJ Abit, AH Ramirez and S Iwakami critically revised the manuscript. All authors read and approved the final manuscript.

## Data Availability Statement

The data that support the findings of this study are available from the corresponding author upon reasonable request.

## References

Bakkali, Y., Ruiz-Santaella, J.P., Osuna, M.D., Wagner, J., Fischer, A.J., De Prado, R., 2007. Late watergrass (*Echinochloa phyllopogon*): Mechanisms involved in the resistance to fenoxaprop-p-ethyl. Journal of Agricultural and Food Chemistry 55(10), 4052–4058. 10.1021/jf0624749

Casimero, M., Abit, M. J., Ramirez, A. H., Dimaano, N. G., Mendoza, J., 2023. Herbicide use history and weed management in Southeast Asia. Advances in Weed Science 40, e020220054. 10.51694/AdvWeedSci/2022;40:seventy-five013

Chauhan, B.S., Johnson, D.E., 2010. Implications of narrow crop row spacing and delayed *Echinochloa colona* and *Echinochloa crus-galli* emergence for weed growth and crop yield loss in aerobic rice. Field Crops Research 117, 177–182. 10.1016/j.fcr.2010.02.014

Damalas, C.A., Dhima, K. V., Eleftherohorinos, I.G., 2008. Morphological and Physiological Variation among Species of the Genus *Echinochloa* in Northern Greece. Weed Science 56, 416–423. 10.1614/ws-07-168.1

Dimaano, N.G., Yamaguchi, T., Fukunishi, K., Tominaga, T., Iwakami, S., 2020. Functional characterization of cytochrome P450 CYP81A subfamily to disclose the pattern of cross-resistance in *Echinochloa phyllopogon*. Plant Molecular Biology 102, 403–416. 10.1007/s11103-019-00954-3

Dimaano, N.G. and Iwakami, S., 2021. Cytochrome P450-mediated herbicide metabolism in plants: current understanding and prospects. Pest Management Science, 77(1), 22–32.

Fertilizer and Pesticide Authority, 2025. List of registered agricultural pesticides. Quezon City: Fertilizer and Pesticide Authority. https://fpa.da.gov.ph/NW/images/FPAfiles/DATA/Regulation/Pesticide/Files-2022/Products/Products-Aug082022.pdf (accessed 30 May 2025)

Gianessi, L., Williams, A., 2012. Increasing Cost of Labor in the Philippines Promotes Herbicide Applications in Rice. International Pesticide Benefits Case Study No, 56.

Hall, T., 2011. BioEdit: An important software for molecular biology Software Review. GERF Bulletin of Biosciences 2(1):60–61.

Heap, I., 2014. Global perspective of herbicide-resistant weeds. Pest Management Science 70(9), 1306–1315. 10.1002/ps.3696

Heap, I., 2025. The International Herbicide-Resistant Weed Database. https://weedscience.org/Home.aspx (accessed 25 September 2025).

Im S. H., P.M.W., Y.M.J., and K.D.S., 2009. Resistance to ACCase inhibitor cyhalofop-butyl in *Echinochloa crus-galli* var. *crus-galli* collected in Seosan, Korea. Korean Journal of Weed Science 29, 178–184.

Iwakami, S., Endo, M., Saika, H., Okuno, J., Nakamura, N., Yokoyama, M., Watanabe, H., Toki, S., Uchino, A. and Inamura, T., 2014. Cytochrome P450 CYP81A12 and CYP81A21 are associated with resistance to two acetolactate synthase inhibitors in *Echinochloa phyllopogon*. Plant Physiology 165(2), 618–629.

Iwakami, S., Hashimoto, M., Matsushima, K. Ichi, Watanabe, H., Hamamura, K., Uchino, A., 2015. Multiple-herbicide resistance in *Echinochloa crus-galli* var. *formosensis*, an allohexaploid weed species, in dry-seeded rice. Pesticide Biochemistry and Physiology 119, 1–8. 10.1016/j.pestbp.2015.02.007

Iwakami, S., Kamidate, Y., Yamaguchi, T., Ishizaka, M., Endo, M., Suda, H., Nagai, K., Sunohara, Y., Toki, S., Uchino, A. and Tominaga, T., 2019. CYP 81A P450s are involved in concomitant cross-resistance to acetolactate synthase and acetyl-CoA carboxylase herbicides in *Echinochloa phyllopogon*. New Phytologist 221 (4), 2112–2122.

Iwakami, S., Ishizawa, H., Sugiura, K., Kashiwagi, K., Oga, T., Niwayama, S., Uchino, A., 2024. Syntenic analysis of ACCase loci and target-site-resistance mutations in cyhalofop-butyl resistant *Echinochloa crus-galli* var. *crus-galli* in Japan. Pest Management Science 80, 627–636. 10.1002/ps.7789

Juliano, L.M., Casimero, M.C., Llewellyn, R., 2010. Multiple herbicide resistance in barnyardgrass (*Echinochloa crus-galli*) in direct-seeded rice in the Philippines. International Journal of Pest Management 56, 299–307. 10.1080/09670874.2010.495795

Lonhienne, T., Garcia, M.D., Low, Y.S., Guddat, L.W., 2022. Herbicides that inhibit acetolactate Synthase. Frontiers of Agricultural Science and Engineering 9, 155–160. 10.15302/J-FASE-2021420

McCourt, J.A., Siew Pang, S., King-Scott, J., Guddat, L.W., Duggleby, R.G., 2006. Herbicide-binding sites revealed in the structure of plant acetohydroxyacid synthase. Proceedings of the National Academy of Sciences 103(3), 569–573. 10.1073/pnas.0508701103

Pan, L., Guo, Q., Wang, J., Shi, L., Yang, X., Zhou, Y., Yu, Q., Bai, L., 2022. CYP81A68 confers metabolic resistance to ALS and ACCase-inhibiting herbicides and its epigenetic regulation in *Echinochloa crus-galli*. Journal of Hazardous Materials 428, 128225. 10.1016/J.JHAZMAT.2022.128225

Powles, S.B., & Yu, Q., 2010. Evolution in Action: Plants Resistant to Herbicides. Annual Review of Plant Biology 61, 317–347. 10.1146/annurev-arplant-042809-112119

R Core Team, 2018. R: a language and environment for statistical computing. R Foundation for Statistical Computing, Vienna, Austria. https://www.r-project.org/ (accessed 2 March 2024)

Ritz, C., Baty, F., Streibig, J.C., Gerhard, D., 2015. Dose-response analysis using R. PLoS One 10. 10.1371/journal.pone.0146021

Saika, H., Horita, J., Taguchi-Shiobara, F., Nonaka, S., Nishizawa-Yokoi, A., Iwakami, S., Hori, K., Matsumoto, T., Tanaka, T., Itoh, T., Yano, M., Kaku, K., Shimizu, T., Toki, S., 2014. A novel rice cytochrome P450 gene, CYP72A31, confers tolerance to acetolactate synthase-inhibiting herbicides in rice and arabidopsis. Plant Physiology 166, 1232–1240. 10.1104/pp.113.231266

Seefeldt, Steven S., Jens Erik Jensen, and E. Patrick Fuerst. 1995. “Log-logistic analysis of herbicide dose-response relationships.” Weed Technology 9:2, 218–227.

Smith, R.J., 1988. Weed thresholds in southern US rice, *Oryza sativa*. Weed Technology 2, 232–241. 10.1017/S0890037X00030505

Suda, H., Kubo, T., Yoshimoto, Y., Tanaka, K., Tanaka, S., Uchino, A., Azuma, S., Hattori, M., Yamaguchi, T., Miyashita, M., Tominaga, T., Iwakami, S., 2023. Transcriptionally linked simultaneous overexpression of P450 genes for broad-spectrum herbicide resistance. Plant Physiology 192, 3017–3029. 10.1093/plphys/kiad286

Tal A., R.M.L., S.G.R., S.A.L., H.J.C., 1993. Glutathione Conjugation: A Detoxification Pathway for Fenoxaprop-ethyl in Barley, Crabgrass, Oat, and Wheat. Pesticide Biochemistry and Physiology 46, 190–199. 10.1006/pest.1993.1050

Tanigaki, S., Uchino, A., Okawa, S., Miura, C., Hamamura, K., Matsuo, M., Yoshino, N., Ueno, N., Toyama, Y., Fukumi, N., Kijima, E., Masuda, T., Shimono, Y., Tominaga, T., Iwakami, S., 2021. Gene expression shapes the patterns of parallel evolution of herbicide resistance in the agricultural weed *Monochoria vaginalis*. New Phytologist 232, 928–940. 10.1111/nph.17624

Tranel, P.J., Wright, T.R., 2002. Resistance of weeds to ALS-inhibiting herbicides: what have we learned? Weed Science 50, 700–712. 10.1614/0043-1745(2002)050[0700:RROWTA]2.0.CO;2

Wu, D., Shen, E., Jiang, B., Feng, Y., Tang, W., Lao, S., Jia, L., Lin, H.Y., Xie, L., Weng, X., Dong, C., Qian, Q., Lin, F., Xu, H., Lu, H., Cutti, L., Chen, H., Deng, S., Guo, L., Chuah, T.S., Song, B.K., Scarabel, L., Qiu, J., Zhu, Q.H., Yu, Q., Timko, M.P., Yamaguchi, H., Merotto, A., Qiu, Y., Olsen, K.M., Fan, L., Ye, C.Y., 2022. Genomic insights into the evolution of *Echinochloa* species as weed and orphan crop. Nature Communications 13(1), 689. 10.1038/s41467-022-28359-9

Ye, C.Y., Wu, D., Mao, L., Jia, L., Qiu, J., Lao, S., Chen, M., Jiang, B., Tang, W., Peng, Q., Pan, L., 2020. The genomes of the allohexaploid *Echinochloa crus-galli* and its progenitors provide insights into polyploidization-driven adaptation. Molecular Plant 13(9), 1298–1310. 10.1016/j.molp.2020.07.001

Yu, Q., Powles, S., 2014. Metabolism-based herbicide resistance and cross-resistance in crop weeds: A threat to herbicide sustainability and global crop production. Plant Physiology 166, 1106–1118. 10.1104/pp.114.242750

Zhou, Q., Liu, W., Zhang, Y., Liu, K.K., 2007. Action mechanisms of acetolactate synthase-inhibiting herbicides. Pesticide Biochemistry and Physiology 89, 89–96. 10.1016/J.PESTBP.2007.04.004

